# Japanese Encephalitis Virus Infected Microglial Cells Secrete Exosomes Containing *let-7a/b* that Facilitate Neuronal Damage via Caspase Activation

**DOI:** 10.1101/324715

**Authors:** Sriparna Mukherjee, Irshad Akbar, Bharti Kumari, Sudhanshu Vrati, Anirban Basu, Arup Banerjee

## Abstract

Extracellular microRNAs (miRNAs) are essential for the cell to cell communication in the healthy and diseased brain. MicroRNAs released from the activated microglia upon neurotropic virus infection may exacerbate CNS damage. Here, we identified *let-7a* and *let-7b* (*let-7a/b*) as the overexpressed miRNAs in Japanese Encephalitis virus (JEV) infected microglia and assessed their role in JEV pathogenesis. We measured the *let-7a/b* expressions in JEV infected post-mortem human brains, mice brains and in mouse microglial N9 cells by the qRT-PCR and *in situ* hybridization assay. The interaction between *let-7a/b* and NOTCH signaling pathway further examined in Toll-like receptor 7 knockdown (TLR7 KD) mice to assess the functions. Exosomes released from JEV infected or let-7a/b mimic transfected N9, and HEK-293 cells were isolated and evaluated their function. We observed an upregulation of *let-7a/b* in the infected brains as well as in microglia. Knockdown of TLR7 or Inhibition of let-7a/b suppressed the JEV induced NOTCH activation possibly via NF-κB dependent manner and subsequently, attenuated JEV induced TNFα production in microglial cells. Further, exosomes secreted from JEV-infected microglial cells specifically contained *let-7a/b*. Exosomes overexpressed with *let-7a/b* were injected into BALB/c mice as well as co-incubated with mouse neuronal (Neuro2a) cells, or primary cortical neuron resulted in caspase activation leading to neuronal damage in the brain. Thus, our results provide evidence for the multifaceted role of *let-7a/b* miRNAs and unravel the exosomes mediated mechanism for JEV induced pathogenesis.

## Introduction

MicroRNAs (miRNAs) are a class of regulatory non-coding RNAs that regulate many biological processes within a cell by direct binding to mRNA, thus influence protein abundance (1). Recent evidence demonstrates that miRNAs can also affect RNA virus replication and pathogenesis through virus-mediated changes in the host transcriptome (2–9). Furthermore, microRNAs have been detected extracellularly and circulating in the blood, CSF and other body fluids (10). These exosomal microRNAs are thought to be released from cells as encapsulated in the vesicles, called as exosomes. These miRNAs containing exosomes can be taken up by specific target cells within the same or in remote tissues where they can exert their repressive function (11, 12). These characteristics make extracellular microRNAs ideal candidates for intercellular communication over short and long distances (13).

In the brain, the exosomes mediated neuron microglia communication recently discussed in few reports (14–16). Most of the cells in the CNS, including neurons, astrocytes, oligodendrocytes, and microglia shed exosomes. These extracellular vesicles are secreted by brain cells under both normal and pathological conditions and transfer specific signal to the neighboring cells. Transfer of miRNAs via exosome may act as a principal mediator for neuroinflammation (17).

Microglia is the brain resident macrophages and plays a vital role in mediating inflammation (18, 19). Neuroinflammation is the hallmark of Japanese Encephalitis virus (JEV) infection. JEV is a neurotropic virus mainly infects children between 1–5 years of age and leads to the permanent neuronal damage, motor deficits, and memory loss (20). In spite of vaccines available against JEV, the case-fatality rate among those with encephalitis can be as high as 30% (http://www.who.int/immunization/diseases/japanese_encephalitis/en/). In last few years, our group and others have shown that JEV infection can cause a significant change in the intracellular and extracellular miRNA profile in the brain and many of the miRNAs possibly play a vital role in controlling viral replication and neuroinflammation (5, 7–9, 21–27). However, it is not clear whether microglia cells derived miRNAs mediated communication involves in developing JEV pathogenicity.

Previously we have identified several significant miRNAs that differentially expressed during JEV infection in the microglial cells (7). Among them, we choose miRNAs *let-7a* and *let-7b* for the further studies because of the following reasons; (a) MiRNAs let-7a/b, are the highly abundant regulator of gene expression in the CNS. (b) They can act as a TLR7 ligand (28) which is also a key regulator of innate immunity against JEV (29). (c) The let-7 family miRNAs are highly conserved across the species (30); (d) these two miRNAs significantly upregulated during JEV infection in human microglial (CHME3) cells (7). In this present study, we explored the role of *let-7a* and *let-7b* in JEV pathogenesis. We demonstrated that upregulation of *let-7a/b* in microglial cells facilitated Proinflammatory cytokine production by interacting with TLR7 and NOTCH pathway. Further, *let-7a/b* released in the extracellular vesicles and accelerated the neuronal damage via activation of caspases.

## Results

### JEV infection induces *let-7a* and *let-7b* expression in microglial cells and infected brain

To understand how JEV infection modulates miRNAs, we previously conducted a global human miRNA array study to identify differentially expressed miRNAs in human microglial cells (7). Of the several miRNAs modulated during infection, we focused here on the miRNAs that belong to the let-7 family. The human let-7 family of miRNA contains 12 members of miRNA.

However, only eight miRNAs of the let-7 family (Let-7a/b/c/d/e/f/g) were found to be modulated during infection. Among them, *let-7a* and *let-7b* upregulated in CHME3 cells at 48 h post infection (hpi), and rests were significantly down-regulated (Fig S1).

As this study focused on *let-7a* and *let-7b*, we validated their expression in JEV infected postmortem human brain; JEV-infected mice brain and in mouse microglial N9 cells by the qRT-PCR assay. Human post-mortem brain tissue sections obtained from the Human Brain Tissue Repository, National Institute of Mental Health and Neurosciences (NIMHANS), Bangalore, India, as described earlier (9). The qPCR data confirmed to the microarray data.

The upregulated expression of *let-7a* and *let-7b* observed in JEV infected human brain samples when compared against mock-infected controls (Fig. 1A). Even in JEV infected mice brains, the results were similar to those observed in JEV-infected human brains (Fig. 1B). Further, we evaluated time-dependent (12 and 24 h) expression of *let-7a/b* in JEV-infected N9 cells. We observed in N9 cells that JEV infection resulted in a time-dependent increase in the *let-7a/b* expression (Fig. 1C). The rise in *let7a/b* level prompted us to further explore the role of *let-7a/b* in development of JEV pathogenesis.

**Fig. 1.**
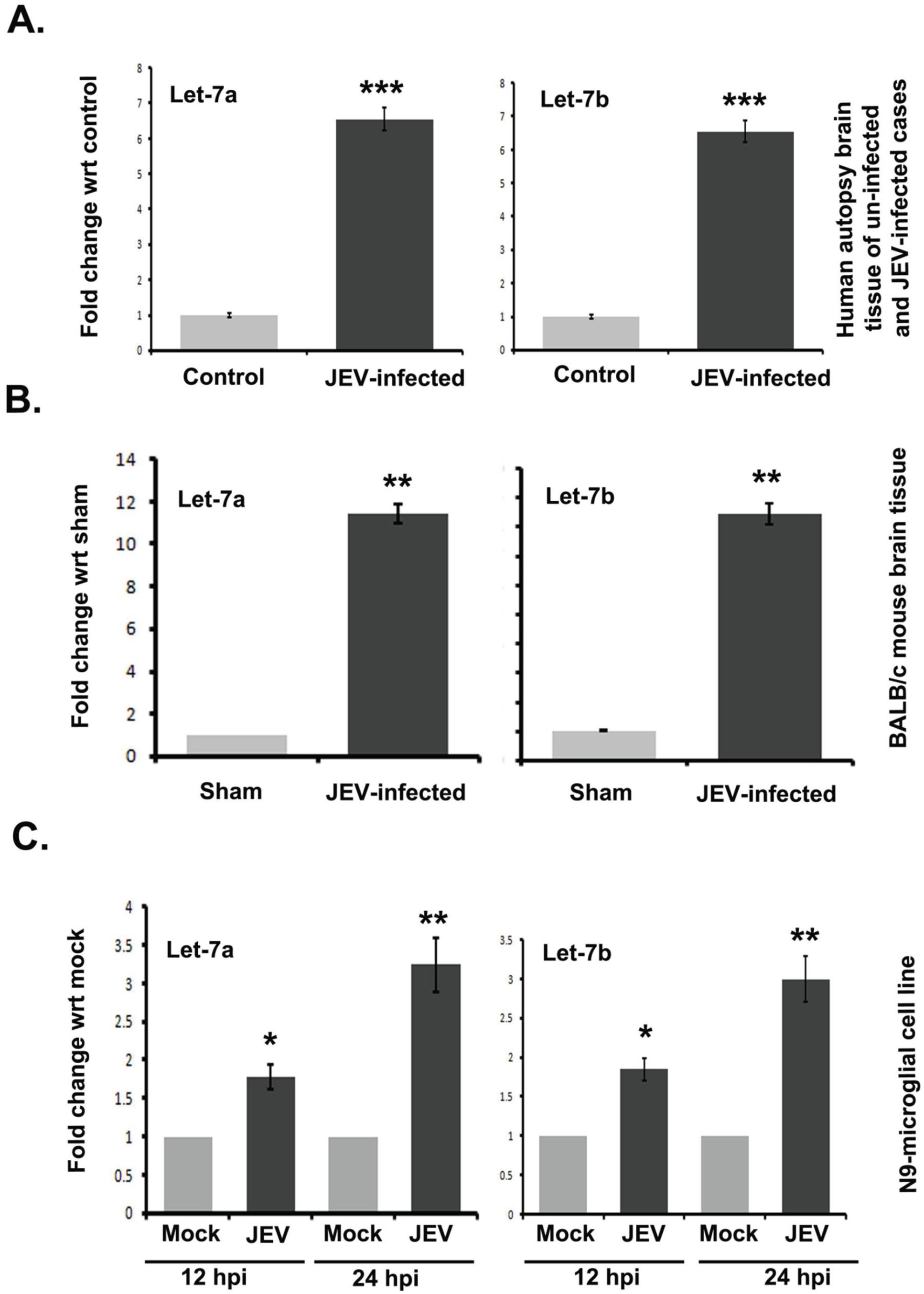
*Let-7a/b* expression is upregulated during JEV infection. **(A)** Graphs showing qRT-PCR results of *let-7a/b* in postmortem JEV-infected and non-JE human brain samples. **(B)** Relative expression of *let-7a/b* in JEV-infected BALB/c mouse brain. BALB/c mice were infected with JEV (3×10^5^ PFU) or mock infected with PBS and brain samples were collected after the 6^th^ day of infection for analysis of *let-7a/b* expression using qRT-PCR. **(C)** Expression of *let-7a/b* in JEV-infected N9 cells. N9 cells were exposed to JEV or mock infected for 12 and 24 h, and expressions of let-7a/b were evaluated using qRT-PCR. * *P*< 0.05 or ** *P*< 0.01 compared to uninfected control. Data are represented as mean ± S.D. from 3 independent experiments.

### Modulation of TLR7 affects JEV induced *let-7a/b* expression in infected mice brain

Due to the presence of repeat **GUUGU** sequence motif in the seed sequence, *let-7a/b* can act as a TLR7 ligand. Our group has already reported TLR7 as one of the critical regulators for JEV pathogenesis (29). Therefore, we explored if TLR7 has any effect on JEV induced *let-7a/b* induction and downstream function. So, we knocked down TLR7 expression in infected mice brain and studied *let-7a/b* expression in wild-type and knocked down brain using qRT-PCR and *in situ* hybridization methods. Treatment of mice with TLR7 specific vivo morpholino significantly reduced TLR7 expression in JEV-infected mice brains as compared to scrambled treated vivo morpholino. We also observed a significant reduction of *let-7a/b* expression in TLR7 knockdown (TLR7 KD) mice brain as compared to both wildtype and scrambled treated JEV infected brain (Fig. 2A). Further, we carried out in situ hybridization for *let-7a/b* in the wild type, and TLR&7 KD JEV infected mice brain section. The purple dots represented the *let-7a/b* signal. As shown in Fig 2B, a strong signal, as well as the increased number of purple spots, observed for *let-7a/b* in wild-type JEV infected mice as compared to mock-infected mice. However, *let-7a/b* positive areas significantly reduced in the TLR7 KD brain (Fig. 2B). Further, we checked phospho-NF-κB (PNF-κB) status in TLR7 KD JEV infected mice brain as *let-7a/b* promoter region contains an NF-κB responsive element (29). Our results suggested that JEV infection induces NF-κB activation in infected mice brain and knockdown of TLR7 reduces the phospho-NF-κB level, suggesting a possible link between NF-κB activation and *let-7a/b* expression (Fig. 2C). These results indicate the TLR7 mediated *let-7a/b* expression, probably dependent on NF-κB activation in JEV infection.

**Fig. 2.**
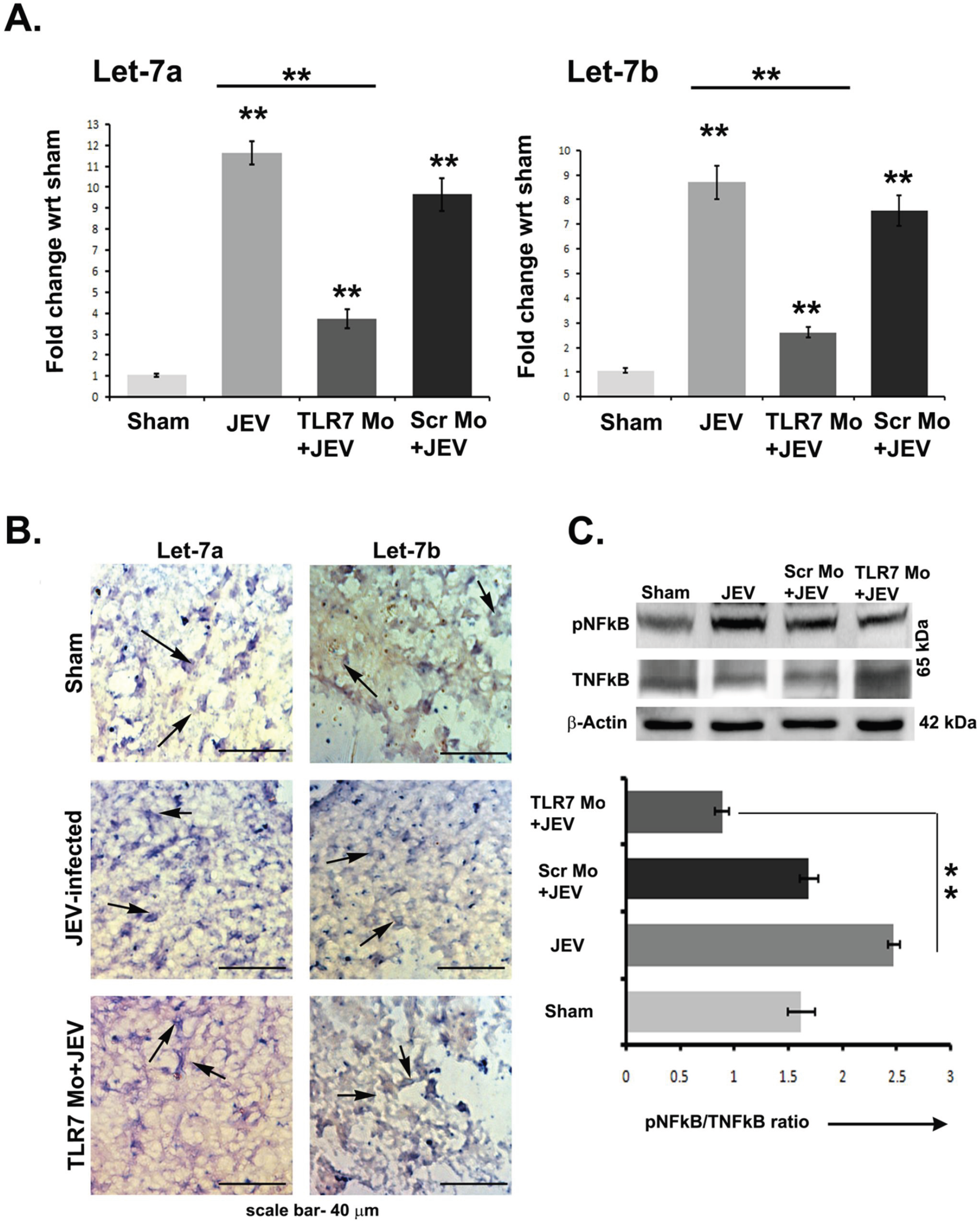
Reduced *let-7a/b* expressions in TLR7 KD BALB/c mice brain. **(A)** Graphs showing qRT-PCR results of *let-7a/b* levels in mice brains belonging to sham, JEV-infected, JEV + Scr-MO and (systemic) JEV + TRL7 KD groups. **(B)** *In situ* hybridization (ISH) showing less abundance of *let-7a* (left panel) and *let-7b* (right panel) in TLR7 KD mice brain as compare to JEV-infected wild-type mice. For ISH study, brain samples were collected seven days post infection (magnification, ×40) and *let-7a/b* detected using specific probes. **(C)** Immunoblots are showing total and phospho-NF-κB levels in mice brains belonging to sham, JEV-infected, JEV + Scr-MO and (systemic/brain) JEV + TRL7 KD groups. The graph (bottom panel) represents densitometry quantification of phospho, and total NF-κB ratio normalized to β-actin.*, P< 0.05 compared to JEV.

### Knockdown of TLR7 or inhibition of *let-7a/b* reduces an active form of NOTCH expression and attenuates JEV induced TNFα expression in microglia

We previously reported NOTCH mediated modulation of inflammation in JEV infected microglial cells (7). Several reports suggested that NOTCH signaling pathway may cross-talk with TLR signaling pathway to control inflammation (31–33). We, therefore, further verified if there is any cross-talk between NOTCH and TLR7 pathway. Upon binding of NOTCH ligand to the receptor, the NOTCH Intracellular Domain (NICD) is cleaved by the gamma-secretase enzyme and transported to the nucleus to activate target genes. We checked NICD level in JEV infected TLR7 KD mice brain. JEV infection induces NICD level in TLR7 wild-type mice brain (Fig. 3A). However, we observed decreased NICD level in TLR7 KD mice brain. Further, using IHC techniques, we checked NICD expression in JEV infected mice brain sections. As shown in Figure 3B, JEV infection significantly upregulated NICD expression (red) in activated microglia, whereas, knockdown of TLR7 significantly reduced NICD expression.

**Fig. 3.**
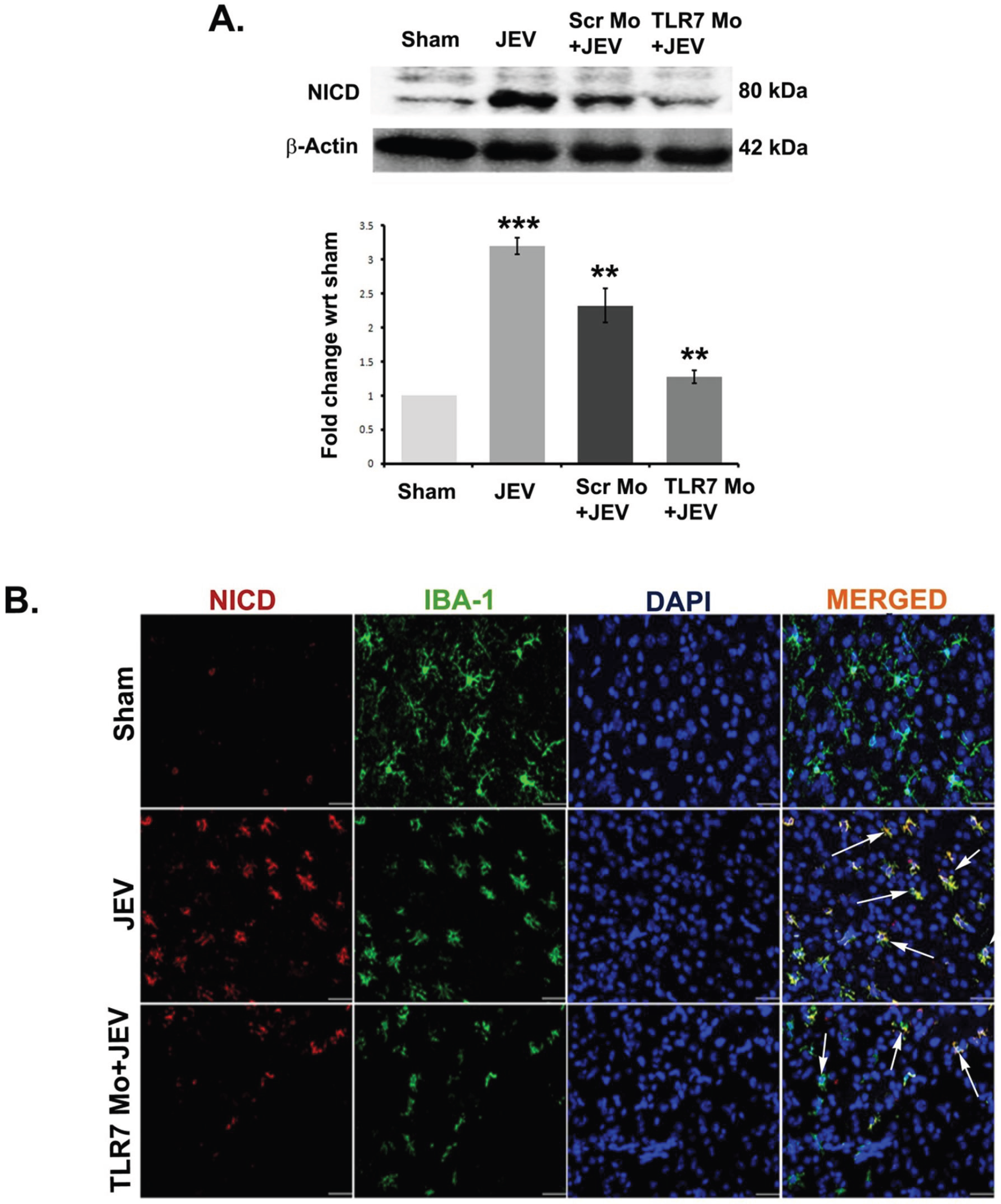
Activated NOTCH expressions reduced in TLR7 KD mice brain upon JEV infection. Immunoblots showing Notch intracellular domain (NICD) levels in mice brains belonging to sham, JEV-infected, JEV + Scr-MO and (systemic/brain) JEV + TRL7 KD groups. The densitometric quantification of NICD normalized against β-actin is shown below. **(B)** Immunohistochemical staining of NICD (red) and activated microglia marker, IBA-1, (green) in brain sections from sham, JEV-infected, JEV + Scr-MO and JEV + TRL7 KD groups. Nucleus stained with DAPI (blue). **, P< 0.005.

We also checked NICD expression in N9 cells. We treated N9 cells with CDPS as CDPS treatment reduced TLR7 expression in N9 cells (Fig. S2A, B). CDPS treatment further reduced JEV induced NF-κB activation (Fig. S2C). We used two different concentrations of CDPS (20 and 30 μM) and infected the cells after twelve hours CDPS treatment. As shown in Figure 4A, JEV infection in N9 cells induces NICD expression. CDPS treatment reduces NICD expression in a dose and time-dependent manner (Fig. 4B). All these data indicate that Notch signaling pathway interacts with the TLR7 pathway.

**Fig. 4.**
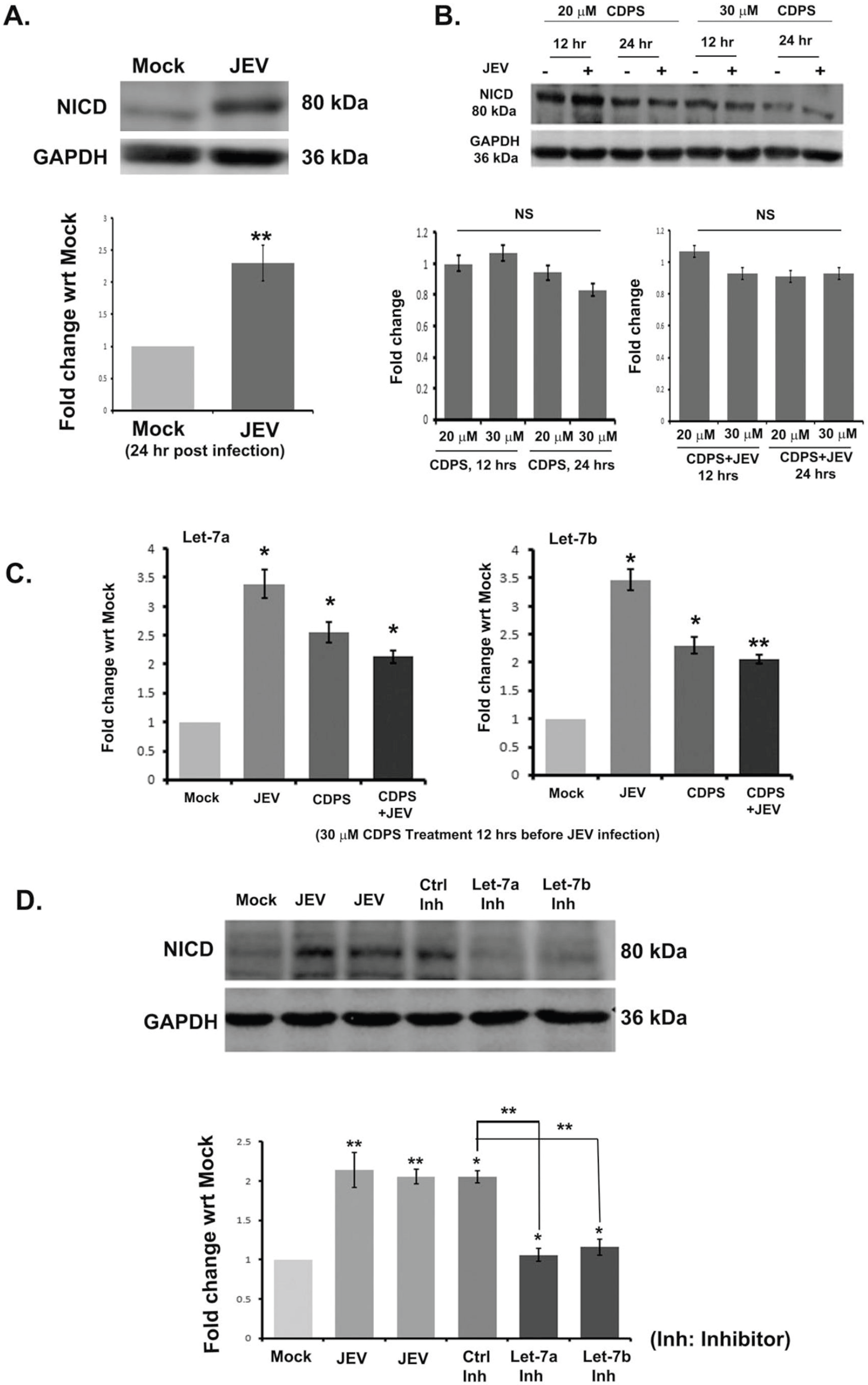
Inhibition of TLR7 function attenuates NOTCH activation. **(A)** Immunoblot represents the higher NICD expression upon JEV infection in N9 cells. Densitometric quantification of NICD band depicted below. **(B)** Relative expression of *let-7a/b* measured in JEV-infected N9 cells upon CDPS (20μM) treatment. **(C)** N9 cells were treated with CDPS (20 and 30 μm) for twelve h, and treated cells were mock-infected or infected with JEV. Cells were lysed at 12, and 24 hours post infection. The NICD expression was studied using Western blot. (D) Expression of NICD in N9 cells either transfected with Control inhibitor or inhibitor specific for *let-7a* or *let-7b* and infected with JEV for 24 h. GAPDH used as loading control. *, P< 0.05, **, P< 0.005.

Further, JEV infection in CDPS treated cells also significantly suppressed *let-7a/b* expression (Fig. 4C). Transfection of *let-7a/b* inhibitor reduced JEV induced NICD expression in N9 cells (Fig. 4D).

NOTCH pathway involves inflammatory cytokine production upon JEV infection (7). *Let-7a/b* expression can induce TNFα in microglia (28). Therefore, to understand the effect of JEV induced *let-7a/b* expression on downstream inflammatory cytokine production, we checked TNFα level (both mRNA and protein) after modulating TLR7 or *let-7a/b* expression. We measured the intracellular TNFα level in CDPS treated N9 cells after 24 hpi by CBA assay. JEV infection in CDPS treated cells significantly suppressed *let-7a/b* expression (Fig. 4C) and also attenuation in TNFα level was observed (Fig. 5A, B). Furthermore, when we treated the N9 cells with inhibitors specific for *let-7a/b*, TNFα level decreased significantly (Fig. 5C).

**Fig. 5.**
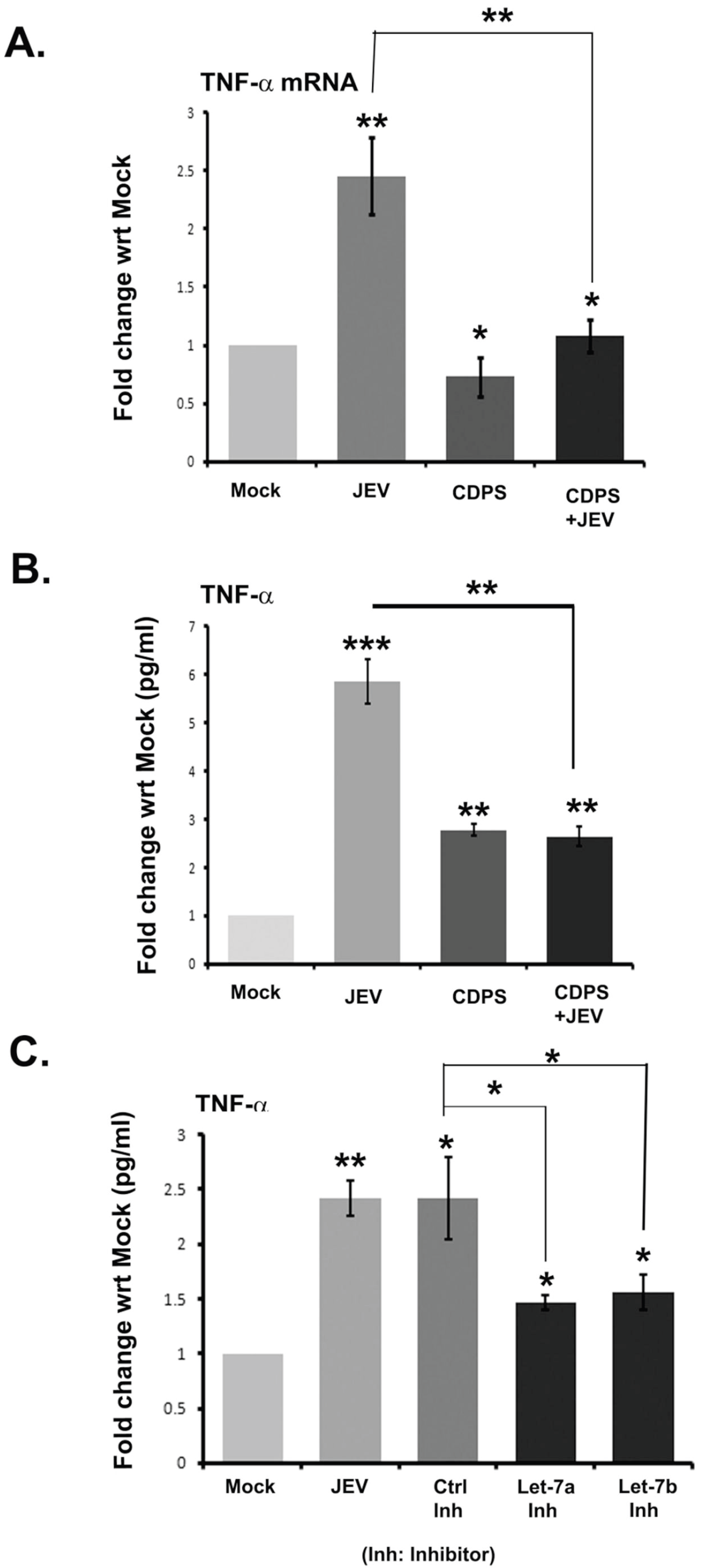
Reduced expressions of TNFα in JEV-infected N9 cells treated with either with CDPS or inhibitor for *let-7a/b.* **(A)** Graph showing qRT-PCR results of TNFα in both CDPS treated and uninfected JEV-infected N9 cells. Glyceraldehyde 3-phosphate dehydrogenase (GAPDH) used as internal control. Data are represented as Mean ± S.D. from 3 independent experiments. * p < 0.05. (B-C) Cytokine Bead Array (CBA) results showing TNFα levels in cell lysates obtained either from CDPS treated JEV infected N9 cells or control inhibitor / let-7a/b inhibitor transfected JEV infected cells. * p < 0.01. Data are represented as mean ± S.D. from 3 independent experiments.

Together these results implicate a possible involvement of let-7-TLR7-NOTCH signaling axis in modulating inflammatory cytokine expression during JEV infection.

### The exosome-mediated release of *let-7a* and *let-7b* in culture supernatant during JEV infection

Since JEV infection induces *let-7a/b* expression in the microglial cells, we checked the level of *let-7a/b* expression in the culture supernatant. For this, we collected the conditioned media after 24 hpi and purified the exosomes. The quality of the purified exosomes was further verified using ZetaSizer, Western blot, and Acetylcholine esterase assay. Our purified exosomes had 100 d.nm in size as depicted in fig. 6A. The expression level of CD63 which is an exosome marker detected in exosomes (Fig. 6B). We also checked the purity of the exosomes by WB against GRP78 (an ER marker) and beta-Actin a cellular protein. The absence of GRP78 and beta-Actin in exosomes confirmed that exosomes were contamination free by cellular breakage or ER burst (Fig. 6B). We also examined the functionality of exosomes by studying acetylcholinesterase activity (Fig. 6C, D). The acetylcholinesterase (AchE) is abundant in exosomes. Exosome purified from JEV infected N9 cells, exhibited increased AchE activity, whereas, exosome released after pretreatment of GW4869, showed significantly reduced AchE activity in a dose-dependent manner, suggesting less exosome production and its release from the cells. We extracted the RNA from the purified exosomes, and by the qRT-PCR approach, we quantified the level of let-7a/b. As compared to mock, exosome derived from JEV infected N9 cells exhibited a higher level of *let-7a/b* expression. Blockade of exosome generation with GW4869 significantly decreased *let-7a/b* level in the culture supernatant (Fig. 6F). However, the intracellular level of *let-7a/b* remained upregulated (Fig. 6E). All these data confirm the release of *let-7a/b* in the circulation through exosomes.

**Fig. 6.**
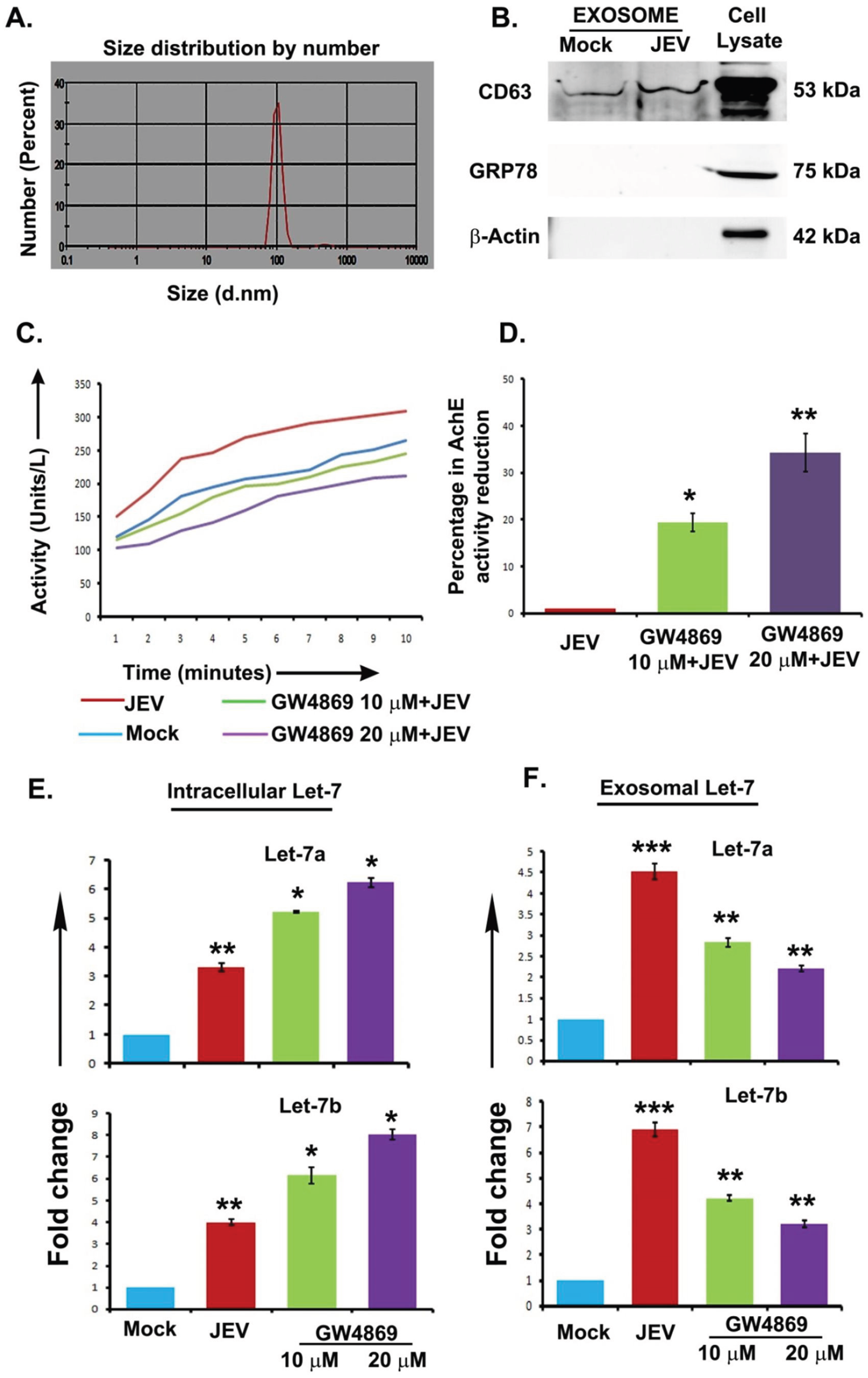
*let-7a/b* released into the culture supernatant via exosome. N9 cells were mock infected or infected with JEV. After 24 hpi, exosomes were purified from cell culture supernatant. **(A)** Graph showing average size of the purified exosomes as measured by ZetaSizer. **(B)** Immunoblot showing CD63, GRP78, Actin expression level in exosomes and whole cell lysate. **(C)** The graph represents functional Acetylcholine Esterase (AchE) activity in exosomes released from mock, JEV infected or pretreatment with exosome biosynthesis inhibitor, GW4869 N9 cells. **(D)** Bar plot showing the percentage of AchE activity reduction upon GW4869 treated N9 cells. **(E-F)** Intracellular and extracellular level of *let-7a/b* in GW4869 treated JEV-infected N9 cells. The graph plotted after normalizing the data against U6 (intracellular) or *cel-miR-39* spike RNA (extracellular) obtained from the qRT-PCR assay. Data are represented as mean ± S.D. from 3 independent experiments.

### Extracellular *let-7a/b* induces neuronal damage via caspase activation

TLR7 is primarily expressed in immune cells but is also present in neurons. Previously our group has shown the critical role of TLR7 expressed in the neuron in mediating JEV pathogenesis (29). Having demonstrated that neurons express TLR7 in our previous study, we tested here the effect of extracellular *let-7a/b* in the brain and on neurons. For this, we first injected mimic overexpressed N9 cells derived exosomes intracranially into the mice brain and collected the mice brain at six days post-injection. The brain cell lysates were subjected to Western blot analysis and checked the expression status of cleaved Poly (ADP-ribose) polymerase (cPARP), cleaved Caspases. As shown in Fig 7A, as compared to control mimic containing exosomes, *let-7a* and *let-7b* containing exosomes treatment caused significant upregulation of cPARP and Caspase 9 in the mice brain. Next, we tested the effect of microglial cell-derived exosomes on neuronal cells. We observed that microglia-derived exosome containing RNAs could be transferred into the neuronal cells by incubating cells with the exosomes (Fig. S3). We have purified the exosomes released in the conditioned media of N9 cells and incubated neuronal (Neuro2a) cells with these exosomes. As compared to uninfected exosomes, JEV infected cells derived exosomes induced more cPARP, Caspase 3, 7 and 9 activations. The additive effect observed with exosome released from mimic transfected cells (Fig. 7B). However, purified exosomes when incubated with freshly isolated mice cortical neurons, Caspase-3 activation was more prominent in the *let-7b* overexpressed exosomes as compared to control mimic expressed exosomes (Fig. 7C, D).

**Fig. 7.**
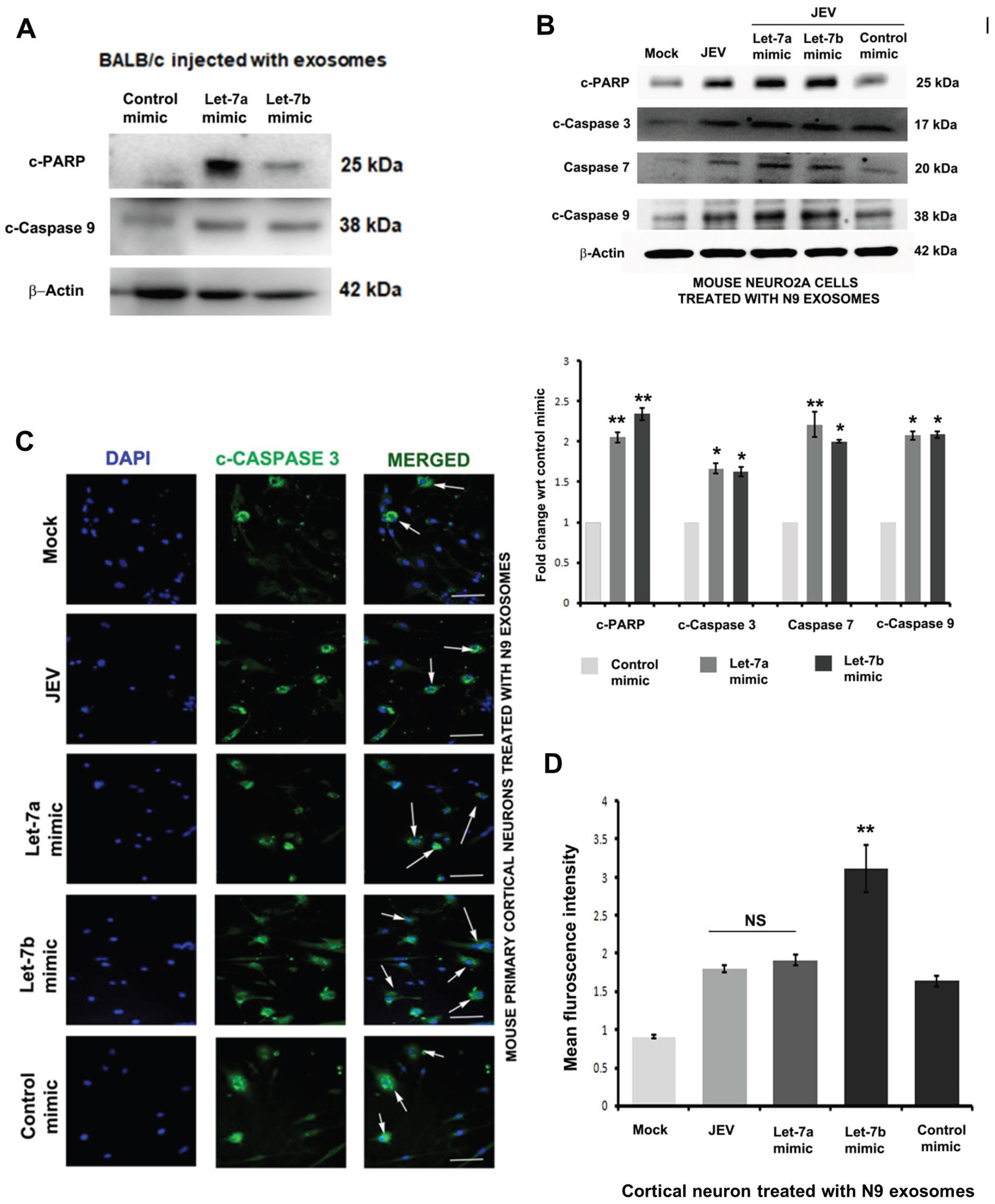
Microglial cell-derived *let-7a/b* containing exosomes induce caspase activation upon incubation with neuronal cells. **(A)** Cleaved PARP and cleaved activated Caspase 9 level in exosome injected whole brain lysate were shown. The β-actin level was used to normalize protein load in each well. **(B) N9** cells were transfected with Control Mimic, Mimic *let-7a* or mimic *let-7b* followed by JEV infection for 24 h. Cells were harvested and examined for cleaved PARP (cPARP), cleaved Caspase 3, Caspase 7 and cleaved Caspase 9 expression by western blot analysis. The graph (below) depicts the densitometry quantification of the relative expression of proteins as compared to control mimic. **(C)** Representative images illustrating activated caspase expression (green) in mouse cortical neurons upon incubation with exosomes released from microglial cells after transfection of control mimic, or mimic specific for *let-7a* and *let-7b* for 24 h. Exosomes purified from mock or JEV infected microglial cells were also incubated with cortical neurons and activated caspase 3 expressing cells were shown. **(D)** Mean fluorescence intensity of three different microscopic fields is summarized in the bar graph. * P< 0.05, ** P< 0.005.

Since *let-7a/b* are present in the exosomes, we hypothesized that transfer of let-7a/b through exosomes could accelerate neuronal damage in the cultured neurons. To confirm the specificity of *let-7* as the causative factor in this context, we further used HEK-293 cells as these cells exhibited a low level of endogenous *let-7a/b* (28, 34). We overexpressed *let-7a/b* in HEK-293 cells by transfecting mimic and purified the exosomes. Freshly prepared exosomes added to neuronal cells. As compared to control mimic treated exosomes, *let-7b* overexpressed exosomes caused significant damage through activation of the PARP, Caspase 3 and Caspase 9. On the other hand, only substantial PARP cleavage was evident in neuronal cells incubated with *let-7a* overexpressed exosomes (Fig. 8A). Further, freshly isolated cortical neurons when incubated with HEK-293 cells derived exosomes; Caspase 3 activation was more prominent in cells incubated with *let-7b* mimic transfected exosomes as compared to control mimic transfected exosomes (Fig. 8B, C). Together, these results confirmed the neurotoxic effect of microglia cell-derived *let-7a/b* containing exosomes in JEV infection. More precisely, irrespective of cellular origin, *let-7b* can induce caspase activation and PARP cleavage in the neuronal cells and causes neuronal damage.

**Fig. 8.**
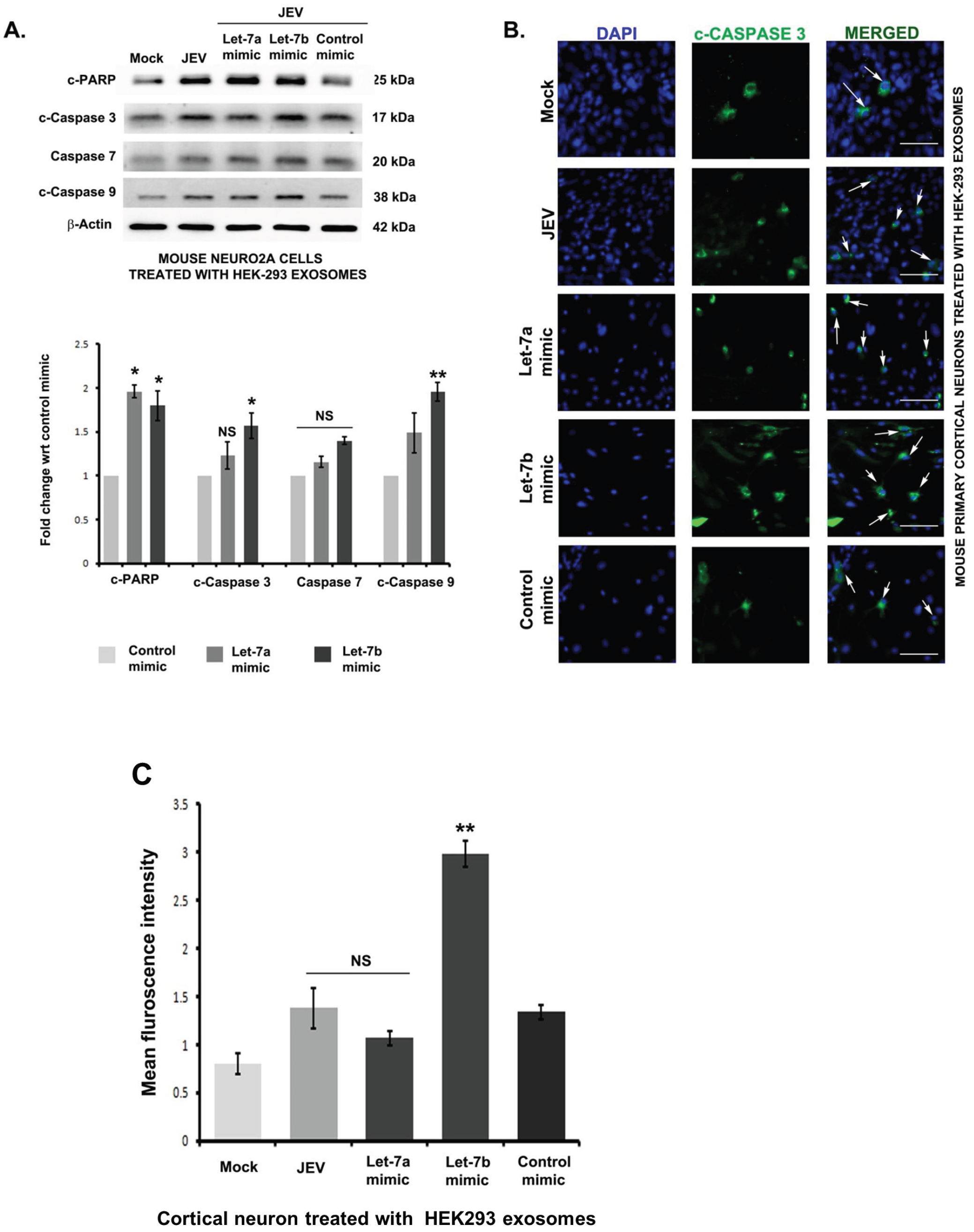
HEK-293 cell-derived *let-7a/b* containing exosomes also induce Caspase activation upon incubation with neuronal cells. **(A)** HEK-293 cells were transfected with Control Mimic, Mimic *let-7a* or mimic *let-7b* followed by JEV infection for 24 h. Cells were harvested and examined for cleaved PARP (cPARPj, cleaved Caspase 3, Caspase 7 and cleaved Caspase 9 expression by Western blot analysis. The graph (below) depicts the densitometry quantification of the relative expression of proteins as compared to control mimic. **(B)** Representative images illustrating activated caspase expression (green) in mouse cortical neurons upon incubation with HEK-293 cells derived exosomes after transfection of control mimic, or mimic specific for *let-7a* and *let-7b* for 24 h. Exosomes purified from mock or JEV infected HEK-293 cells were also incubated with cortical neurons and activated Caspase 3 expressing cells were shown. **(C)** Mean fluorescence intensity of three different microscopic fields is summarized in the bar graph. * P< 0.05, ** P< 0.005.

## Discussion

Cell to cell communication within the brain is essential in maintaining homeostasis, and brain function. The small membrane-bound vesicle called exosomes play a vital role in cell to cell communication by transferring the biologically active component to the distant/neighboring cells. In this study, we have shown that JEV induces *let-7a/b* expression in the microglial cells, facilitates microglia-mediated inflammatory response via interacting with NOTCH signaling pathway. Simultaneously, *let-7a/b* can be packaged into the exosomes, and able to transfer the materials to the neurons where it induces caspase activation and causes neuronal damage.

Let-7 was the first miRNA family discovered in human and highly conserved across species in sequence and function, Misregulation of *let-7a/b* can lead to impair brain function. In our early studies, we observed an upregulation of *let-7a/b* miRNAs in human microglial cells during JEV infection. Since *let-7a* and *let-7b* are highly abundant in brain, changes in the *let-7a/b* level due to JEV infection may associate with pathogenesis. Interestingly, *Let-7a/b* miRNAs may act as a TLR7 ligand and signaling mediated through TLR7 is critical for regulating innate immune response in JEV infection (29). In this present study, we observed that *let-7a/b* expression in JEV infected mice probably interact with TLR7 and modulated JEV induced Notch activation. Notch receptor-mediated signaling is essential in mediating inflammatory response by microglia (35) increases proinflammatory cytokine production during JEV infection (7). As reported earlier, the let-7a/b mediated function can act through Notch and TLR7 signaling pathway (28, 36). Our study further demonstrated a link between *let-7a/b,* TLR7, and Notch in regulating JEV induced TNFα production in microglial cells.

We also observed the reduced level of *let-7a/b* expression upon JEV infection in TLR7 KD mice brain. One of the reasons may be that TLR7 KD reduces the p-NF-κB level in mice brain since *let-7* promoter contains NF-κB responsive elements, one may argue that reduced p-NF-κB expression in TLR7 KD mice brain may indirectly affect *let-7* expression. However, controversial reports regarding the involvement of NF-κB in the *let-7* expression are also evident. Lliopoulos and colleagues found an inverse correlation between NF-κB activation and *let-7* levels (37), Whereas Garzon and colleagues reported NF-κB mediated *let-7* expression during granulocytic differentiation (38). These suggest that the final biological outcome of NF-κB activation on a *let-7* level may vary depending upon the cellular context.

JEV infection causes irreversible damage to neurons. Microglial activation can lead to bystander damage to the neuronal cells possibly through releasing cytokines and chemokines. However, with the discovery of exosome, it is speculated that exosome-mediated transfer of genetic material may affect recipient cell fate.

Our data suggested that *let-7a/b* released in the culture supernatant via exosome. *In vitro* study also confirmed that exosome containing RNA could transfer into the neuronal cells indicating that *let-7a/b* might play a role in cell to cell communication. We observed that microglial cell-derived exosomes upon incubation with neuronal cells, or directly injected into mice brain induced caspase activation. To confirm that *let-7a/b* may be the causative agent and not the other proteins or miRNAs released from infected microglia, we used HEK-293 cells derived exosomes as this cell expressed a low level of *let-7a/b.* Both *let-7a* and *let-7b* overexpressed exosomes (derived from N9 microglia and HEK-293 cells) can induce PARP inactivation in neuronal cells. PARP inactivation occurs in cells where DNA damage is extensive. PARP inactivation occurs due to cleavage by caspases. However, there is a difference in caspase activation pattern in primary neurons when exposed to exosomes. The *let-7b* containing exosomes (derived from N9 and HEK-293 cells) caused significant caspase 3 activations in compare to *let-7a* containing exosomes. These further indicate that transfer of *let-7b* to neuronal cells can accelerate neuronal damage in the infected brain, irrespective of its cellular origin. The PARP cleavage and caspase activation together probably indicate mitochondrial dysfunction caused by *let-7a/b* leading to neuronal damage.

Together with our results suggest that JEV induced *let-7a/b* expression manifested a multifaceted role in JEV pathogenesis. In microglial cells, *let-7a* and *let-7b* can activate NOTCH through cross-talking with TLR7 mediated signaling pathway and induces inflammatory cytokines production. On the other hand, extracellular release of *let-7a/b* via exosomes can transfer to neurons where it can produce neurotoxic effect via activation of caspases. This study unravels the novel exosomes mediated mechanism for virus-induced pathogenesis.

## Methods & Materials

### Ethics statement

Animal experiments were approved by National Brain Research Centre Animal Ethics Committee (Approval no: NBRC/IAEC/2014/96). Animals were maintained according to the guidelines of the Committee for Control and Supervision of Experiments on Animals, Ministry of Environment and Forestry, Government of India. They were kept under a 12 h light/dark cycle with constant temperature and humidity. Food and water supply was *ad libitum*.

### Virus propagation

Suckling BALB/c mice of either sex were intracranially infected with JEV GP78 strain and housed with their mothers. 72-96 h. post-infection, animals developed encephalitic symptoms. At this time, brain samples were collected, and 10% of tissue suspension was prepared in MEM (minimum essential medium) followed by centrifugation at 10000Xg for 30 minutes. The cell culture supernatant was passed through a 0.22 μm sterile filter. Titration of the isolated virus particles was done in porcine kidney (PS) cells by plaque assay.

### Knocking down TLR7 in BALB/c mice brains

TLR7 vivo morpholino was procured from Gene Tools (USA). Knocking down of TLR7 in mouse brain was performed according to the previously published protocol (29). Briefly, ten-day-old BALB/c mice pups of either sex were randomly divided into four groups (Sham, JEV, TLR^KD^+JEV and Scr Mo+ JEV). Mice belonging to TLR KD group received one single intracranial 12.5 mg/kg dose of TLR7 vivo morpholino (5′ TCC GTG TCC ACA TCG AAA ACA CCA T 3′). Simultaneously Scr Mo+ JEV group received one single intracranial dose of Scr morpholino (5′ GAT AAT TCT GGT TTT AAA TTC 3′). Animals were infected with 3×10^5^ PFU of JEV after 48 h. Sham group received the same volume of sterile PBS.

### Cell culture

Mouse neuroblastoma (neuro2a) cells were grown in Dulbecco’s Modified Eagle Medium (DMEM) containing 10% fetal bovine serum. Mouse microglial cell (N9) (a kind gift from Prof. Maria Pedroso de Lima, Centre for Neuroscience and Cell Biology, University of Coimbra, Portugal) was grown in RPMI-1640 medium supplemented with 10% fetal bovine serum. HEK-293 cells (provided by A. Krishnan, Institute of Molecular Medicine, and New Delhi, India) were also cultured in Dulbecco’s Modified Eagle Medium (DMEM) containing 10% fetal bovine serum. All culture media were supplemented with 100 units/ml penicillin-streptomycin.

### Primary cortical neuron culture

Two-day-old BALB/c mouse pups were decapitated under sterile conditions and cortex was isolated in calcium-magnesium free Tyrode solution under a dissecting microscope. Cortex was digested with Trypsin and DNase enzymes to make single cell suspensions. The suspension was passed through a 127 μm pore size nylon mesh (Sefar) to eliminate cellular debris. The filtrate was centrifuged at 1000 rpm to get the cell pellet. Cells were counted in a hemocytometer, and an equal number of cells were seeded in poly-D-lysine (Sigma, USA) coated plates in a neurobasal medium containing 2mM L-glutamine, 1% glucose, 5% FBS, Horse serum and penicillin-streptomycin. After two days, serum was removed from the media to inhibit glial growth and Neuro2, and B27 supplements were added in the cell culture medium. Cells were treated with 20 μM arabinoside one day before viral infection to eliminate glial presence in the culture,

### JEV infection *in vivo* and *in vitro*

JEV, TLR^KD^+JEV and Scr Mo+ JEV group of animals received 3×10^5^ PFU of JEV (GP78) intraperitoneally. Sham group received an equal volume of PBS. Day 5 post-infection onwards animals developed encephalitic symptoms and brain samples were collected after perfusion with cold 1X sterile PBS prepared using DEPC treated water. Samples thus obtained were used in protein, RNA or miRNA isolation. For cryo-sectioning and Immune-histochemistry, perfused brains were kept in 4% PFA for fixation followed by 30% sucrose treatment. All reagents were prepared in DEPC water when brain samples were collected for in-situ hybridization.

N9 cells were grown till 70-80 % confluency followed by serum-free RPMI addition for two h. Cells were treated with 5 MOI of JEV for two h. Then cells were washed for the removal of uninternalized virus and kept in fresh RPMI media for 24 h. After that, cells were harvested for protein or RNA or miRNA isolation.

### Chloroquine treatment in cells

N9 cells were grown till 70% confluency followed by serum-free media addition. Cells were divided into four groups (mock, JEV, only CDPS and CDPS+JEV). Cells were treated with 20 or 30 μM CDPS (Chloroquine diphosphate salt, Sigma, USA) 12 h. before JEV infection. Then cells were washed thoroughly with sterile PBS and infected with 5MOI of JEV for 24 h. Cells of the mock group were treated with PBS.

### Western-Blot

Both tissue and cell samples were lysed using RIPA buffer supplemented with 0.2% sodium orthovanadate and protease inhibitor cocktail (Sigma, USA). Protein concentration was measured using BCA reagent (Sigma, USA). 30 μg of protein samples were separated in SDS-PAGE followed by transfer onto a nitrocellulose membrane. The membrane was blocked using 10% skimmed milk and incubated with primary antibodies overnight at 4^0^C. Suitable secondary antibodies from Vector Laboratories, USA were used after PBST washes on the next day. Blot development was performed in UNITECH imaging system (Cambridge) using ECL reagent from Millipore, CA USA. All the antibodies used for this study were listed in Table S1.

### Immuno-histochemistry

20 μm sections of mouse brain were treated with antigen unmasking solution (Vector Laboratories, USA) at 70°C followed by PBS washes. Sections were blocked with 10% animal serum for one hour and 30 min followed by overnight incubation with an antibody against activated Notch (Abcam, USA, #ab8925) at 4^0^C. Next day, sections were washed with PBS followed by one h incubation with secondary rabbit polyclonal antibody (fluorochrome-conjugated) at room temperature. After extensive washes, brain sections were mounted in DAPI (Vector Laboratories, USA) and observed under fluorescence microscope (Zeiss, Germany).

### Cytokine bead array

TNFα concentration in CDPS treated N9 cells were measured by flow cytometry using a mouse inflammation CBA kit (BD Biosciences, San Diego, CA, USA) as per the manufacturer’s instructions. Data were obtained in FACS Calibur (Becton Dickinson, San Diego, CA, USA).

### MicroRNA isolation and qPCR analysis

MicroRNA isolation from both mouse brain and N9 cells were performed using miRNeasy Mini Kit (Qiagen, USA) following the manufacturer’s instructions. miScript II RT kit (Qiagen) was used for cDNA synthesis. Reaction conditions were: 37^0^C for 60 min and 95^0^C for 5 min. For qPCR reactions, miScript SYBR green PCR kit (Qiagen) was used. Reaction conditions were: 95^0^C for 15 min(1 cycle), 40 cycles at 94^0^C for 15 sec, 55^0^C for 30 sec and 70^0^C for 30 sec. SNORD-68 or U6 were used as internal controls, and qPCR data were analyzed using comparative delta CT(2^-ΔΔCT) method. The miRNA isolation from exosomes was done by the same method as described. Synthetic spiked *C. elegans* miR-39 was added to the exosomes before miRNA isolation, and this was used as endogenous control.

### MicroRNA isolation from human brain sections

miRNA was isolated from paraffin-embedded basal ganglia region of JEV infected human autopsy tissue (CSF positive for JEV −IgM). Age-matched non-JEV control samples were accidental cases with least possible trauma to the brain. Tissue samples were collected from Human Brain Bank, NIMHANS, Bangalore according to institutional ethics and confidentiality of the subjects. miRNA was extracted using miRNeasy FFPE Kit (Qiagen) following manufacturer’s protocols. 20μm sections were deparaffinized using xylene treated with proteinase K and processed for miRNA isolation. The cDNA was prepared using miScript II RT kit (Qiagen) as stated before.

### Transfection of cells with miRNA mimic and inhibitor

N9 and HEK-293 cells were transfected with miRCURY LNA let-7a/7b-5p mimic or miRCURY LNA *let-7a/7b-5p* inhibitor to overexpress or downregulate *let-7a/7b* using the HiPerfect Transfection Reagent (Qiagen) following manufacturer’s instructions. 24 h post-transfection, cells were harvested to check for transfection efficiency. Negative control of both mimic and inhibitors (Ambion) was used in each experiment.

### In situ hybridization

20μm cryosections of mouse brain were taken for ISH experiments using miRCURY LNA miRNA ISH optimization kit (Exiqon) as used previously (Menaka paper). Sections were kept in 30% formalin overnight. After PBS wash, sections were treated with 7mg/ml proteinase K at 37^0^C for 15 min. Then, sections were dehydrated using graded ethanol followed by incubation with hybridization buffer containing 50nM *let-7a* (5’-3/56-FAM/AACTATACAACCTACTACCTCA) and *let-7b* (5’-3’/56-FAM/AACCACACAACCTACTACCTCA) probes (Exiqon) for one hr at 55^0^C in a humid chamber. Later, sections were washed in 5×, 1× and 0.2 × saline-sodium citrate buffer and blocked in BSA and sheep serum for 15 minutes at room temperature. 5 U/ml anti-Fluorescein-AP Fab fragments (Roche Diagnostics, Germany) were used to cover the sections for one h at room temperature following two PBST (Tween 20) washes. Freshly prepared AP substrate (Roche) was added to the sections for two h at 30^0^C in a humid chamber. AP stop solution (KTBT buffer containing 50 mM Tris-HCl, 150 mM NaCl and ten mM KCl) was used to terminate the reaction. Sections were washed in DEPC water followed by dehydration using graded alcohol. They were mounted using DPX reagent (Qualigens). Sections were observed in a Leica DMRXA2 microscope.

### Isolation of Exosomes from cell culture supernatant

Exosomes from N9 and HEK-293 cell culture supernatant were isolated using total Exosome isolation reagent (Invitrogen) using manufacturer’s instructions. For exosome isolation, virus-infected or mimic transfected cells were washed five times with PBS and then supplemented with serum-free media and incubated for 24 hours. The culture supernatant was collected and centrifuged at 2000Xg for 30 min to remove cell debris. Then, the culture media was transferred to a fresh tube, and 0.5 volumes of isolation reagent were added to that and left overnight at 4^0^C. Next day supernatant was centrifuged at 10,000Xg for one h at 4^°^C. Exosomes pellet was resuspended in the re-suspension buffer provided with the Kit for treatment into cells, and used in miRNA isolation.

### Labeling of Exosomes and treatment into cells and mice brain

Exosomes isolated from N9 cells were labeled using ExoGlow reagent (SBI Biosystems) following manufacturer’s instructions. 100 μl of labeled exosomes were used for 10^5 cells. Exosome isolated from N9 cells were labeled accordingly and added to Neuro2a cells for different time points. Then cells were washed and fixed with 4% PFA for four h at 4^0^C. Entry of exosomes inside the cells was monitored by a counterstain of transferrin receptor (membrane protein) and DAPI and visualized in a Zeiss Apotome microscope (Zeiss, Germany).

Unlabeled N9 exosomes were applied in Neuro2a cells, and primary cortical neurons at a concentration of **30** μg/ml for 24 h and cells were harvested for protein isolation and western-blot.

Ten-day-old BALB/c mouse pups of either sex were randomly divided into five groups. Animals in each group received 70 μg intracranial injections of exosomes isolated from N9 cells (Untreated, JEV, let-7a mimic+ JEV, let-7b mimic + JEV, control mimic + JEV). N9 cells were infected with 5 MOI of JEV and post 24 h of infection; supernatant from the mentioned groups was collected for exosome isolation. Brain samples from the exosome treated animals were collected after six days post administration through 1X sterile PBS perfusion. Collected brain tissues were used for protein isolation and western-blot.

### Immuno-cytochemistry

Freshly isolated mouse primary cortical neurons were treated with exosomes isolated from N9 or HEK-293 cells at a concentration of 30ug/ml for 24 h. The cells were thoroughly washed with PBS, fixed in 4% PFA followed by permeabilization with 0.1% Triton X containing PBS and blocking with animal serum in which the secondary antibody is raised in. The cells were then incubated with primary antibody against Caspase 3 at 4°C overnight [dilution used for primary antibody was 1:200]. The cells were washed with 0.1% Triton X containing PBS and incubated with the fluorochrome-conjugated secondary antibody (1:500) for 1h at room temperature. The cells were then mounted with DAPI (Vector Laboratories, USA) following five washes and were subsequently observed under a fluorescence microscope (Zeiss, Germany).

### Statistical analysis

Data were represented as mean±SD of triplicate experimental sets (n=3). Statistical significance was evaluated using Student’s *t*-test or one-way analysis of variance (ANOVA) followed by Holm-Sidak post hoc test. P-Value < 0.05 was considered to be statistically significant.

## Acknowledgments

Authors sincerely acknowledge Prof. Anita Mahadevan and Prof. SK Shankar, Department of Neuropathology, National Institute for Mental Health and Neurosciences (NIMHANS), Bangalore, for providing autopsied human tissue.

## Author contribution

Conceptualization and Experimental design: Sriparna Mukherjee, Arup Banerjee, Anirban Basu, and Sudhanshu Vrati

The investigation, validation, data curation, and visualization: Sriparna Mukherjee, Irshad Akbar, Bharti Kumari

Supervision and manuscript drafting: Arup Banerjee, Anirban Basu, Sudhanshu Vrati Funding

## Abbreviations

CDPS: Chloroquine diphosphate salt; KD: Knockdown; NICD: Notch intracellular domain; cPARP: cleaved Poly (ADP-ribose) polymerase;

